# Transcriptomic and Proteomic analysis of clear cell foci (CCF) in the human non-cirrhotic liver identifies several differentially expressed genes and proteins with functions in cancer cell biology and glycogen metabolism

**DOI:** 10.1101/2020.02.10.941211

**Authors:** Christoph Metzendorf, Katharina Wineberger, Jenny Rausch, Antonio Cigliano, Diego F. Calvisi, Kristin Peters, Baodong Sun, Daniela Mennerich, Thomas Kietzmann, Frank Dombrowski, Silvia Ribback

## Abstract

Clear cell foci (CCF) of the liver are considered to be pre-neoplastic lesions of hepatocellular adenomas and carcinomas. They are hallmarked by glycogen-overload and activation of AKT/mTOR-signaling.

Here we report the transcriptome and proteome of CCF extracted from human liver biopsies by laser capture microdissection. We mainly found genes and proteins with lower expression in CCF compared to control samples. Of 14 differentially expressed transcripts only 3 were overexpressed and belong to previously uncharacterized long non-coding RNAs. At the protein level, 3 proteins were at least 1.5-fold higher expressed in CCF than in control samples, while 22 proteins were at least 1.5-fold less abundant. Using immunohistochemistry, the reduced expressions of STBD1, USP28, monad/WDR92, CYB5B and HSPE1 were validated in CCF. Knockout of *STBD1*, the gene coding for Starch-binding domain-containing protein 1, in mice did not have an effect on liver glycogen levels. Similiarly, *Usp28* knockout in mice did not affect glycogen storage in diethylnitrosamine-induced liver carcinoma. Thus, these data indicate that reduced expression of STBD1 or USP28 in CCF is unlikely to explain glycogen overload.

Our study revealed several novel proteins and RNAs that are differentially expressed in CCF and have functions relevant for carcinogenesis or the glycogen metabolism.

## INTRODUCTION

The early processes underlying human hepatocellular carcinogenesis are poorly understood. Very diverse conditions, such as the cirrhotic liver, non-cirrhotic liver with glycogen storage disease type I [1] as well as metabolic disorders (alpha-1-antitrypsin deficiency and hemochromatosis) or like obesity [2], hyperinsulinism, alcohol abuse, and type II diabetes mellitus [3,4] are known to be risk factors for hepatocellular carcinoma (HCC) development. While high grade dysplastic nodules in liver cirrhosis are accepted as pre-neoplastic lesions of HCC [5], the situation in the absence of liver cirrhosis is less clear, even though 15-20% of HCC occur in non-cirrhotic livers [6].

To better understand the mechanisms underlying carcinogenesis it is important to better characterize the precursor stages - pre-neoplastic lesions, for improving early diagnosis and treatment of HCC. This becomes more and more important as primary liver cancer is the fifth most frequent malignancy worldwide and the proportion of HCCs in the background of type 2 diabetes and obesity becomes more common [2].

In the human cirrhotic liver, different types of foci of altered hepatocytes were described by Bannasch: glycogen storing foci (clear cell foci, CCF, with pale HE staining), mixed cell foci and basophilic foci [7]. These foci are also well known in diverse animal models of hepatocarcinogenesis [8] and their progression to hepatocellular adenomas and HCC is well described [7,9-11]. Moreover, using the intraportal pancreatic islet transplantation model of hepatocarcinogenesis [12-15], we found the AKT (v-akt murine thymoma viral oncogene homolog)/mTOR (mammalian target of rapamycin) and the Ras (rat sarcoma)/MAPK (mitogen activated protein kinase) pathways activated throughout the development of CCF to HCC, where they play important roles as major oncogenic downstream effectors of insulin signaling [16,17]. The lipogenic phenotype is characterized by increased lipogenesis and storage of lipid droplets. These alterations have also been described in human HCC, where they are associated with unfavorable prognosis [18,19]. Recently, we described that CCF in human non-cirrhotic livers reveal many molecular and metabolic characteristics like pre-neoplastic liver foci of the hormonal model of hepatocarcinogenesis after intraportal pancreatic islet transplantation [20]. Specifically, we found an increased glycogen storage, reduced glucose-6-phosphatase activity and upregulated enzymes of glycolysis and *de novo* lipogenesis, beta-oxidation as well as overexpression of the insulin receptor and activated AKT/mTOR and Ras/MAPK pathways in CCF from both human livers and the rat model [20]. Similiarly, in the mouse, hepatocarcinogenesis is associated with activation of the insulin/AKT/mTOR signaling pathway, the transcriptional regulator ChREBP [16,21], as well as the lipogenic pathway [18,22,23].

Although these data hint to several pathways and regulators, a comprehensive inventory of gene and protein expression in human CCF is missing.

In the current work, we applied microarray analysis and proteomics after laser microdissection as unbiased approaches to identify and to further characterize CCF of human non-cirrhotic liver parenchyma. To this end we compared RNA and protein expression in CCF with neighboring tissue as control and found few genes and proteins with significantly changed expression. Starch-binding domain-containing protein 1 (STBD1), ubiquitin specific peptidase 28 (USP28), WD repeat-containing protein 92 (WDR92)/Monad and heat shock protein family E (Hsp10) member 1 (HSP10) were among the candidates with highest differential expression and their expression was validated by immunohistochemistry.

## RESULTS

### More RNAs have reduced expression in CCF compared to controls

After standard processing of the microarray dataset, as described in materials and methods, we performed the following tests to identify any problems with sample quality, normalization or signal quality. First, we determined how many transcripts were above the non-detection threshold in each sample and could not find any samples to drop out noticeably (Supplementary Figure S1 A). Also, the mean number of transcripts of control and CCF samples did not show any statistically significant difference (Supplementary Figure S1 A and B; n = 18, students t-test, p (unpaired) = 0.104; p (paired) = 0.063). The distributions of log-transformed signal intensities per sample were quite similar between samples (Supplementary Figure S2). From these observations we concluded that the quality of the data was acceptable for further analysis.

Using cluster analysis, samples clustered in a patient-dependent manner in most cases (Figure 1 A), indicating that differences between patients were more extensive than those between CCF and control samples. A larger degree of heterogeneity is not uncommon for human tissue samples, in general, and we suggest, that transcripts with statistically significant differences between CCF and control samples will be quite robust. On the other hand, we may miss differentially expressed genes due to the higher degree of noise in the data. To take the inter-patient heterogeneity into account, we compared RNA expression between CCF and controls by calculating fold-changes (CCF/control) per patient. 14 transcripts (Table 1 and Figure 1 B) had at least 2-fold higher or lower expression in CCF than in control samples, three of these with higher and 11 with lower expression in CCF-samples. Interestingly, all three transcripts with increased expression in CCF coded for long non-coding RNAs (lnc-FOXG1-6:17, LINC01124:6 and LINC02290:27). However, nothing is known about the function of any of these lncRNAs. LINC01124:6 is annotated as a bidirectional, 2129 bp long lncRNA encoded by one exon, while LNC02290:27 is an intergenic lncRNA of 463 bp length encoded on 4 exons (LNCIPEDIA v 5.2, www.lncipedia.org).

**Table 1:**
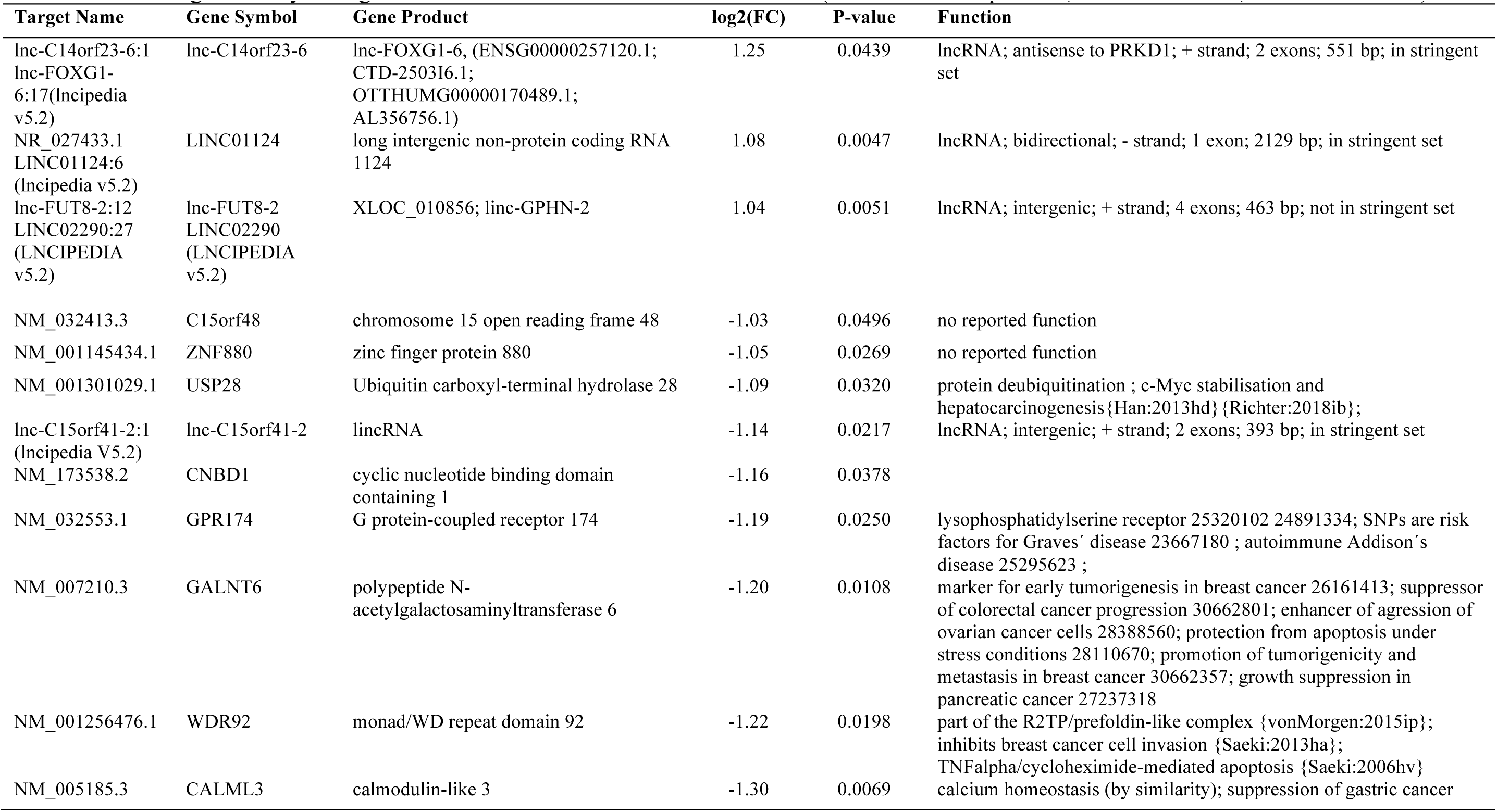
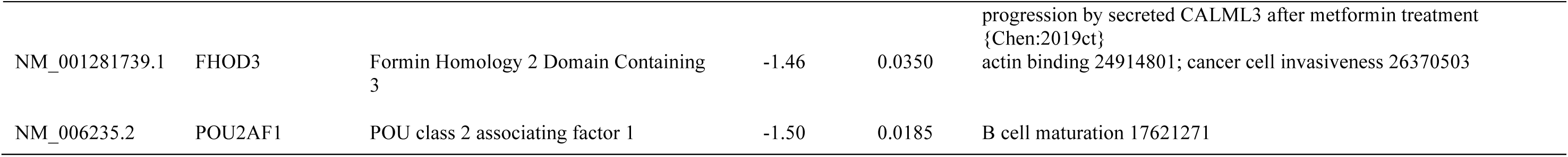
Genes significantly changed between CCF and unaltered liver tissue (at least 2 fold up/down, CCF vs control; P-value <= 0.05)

**Figure 1:**
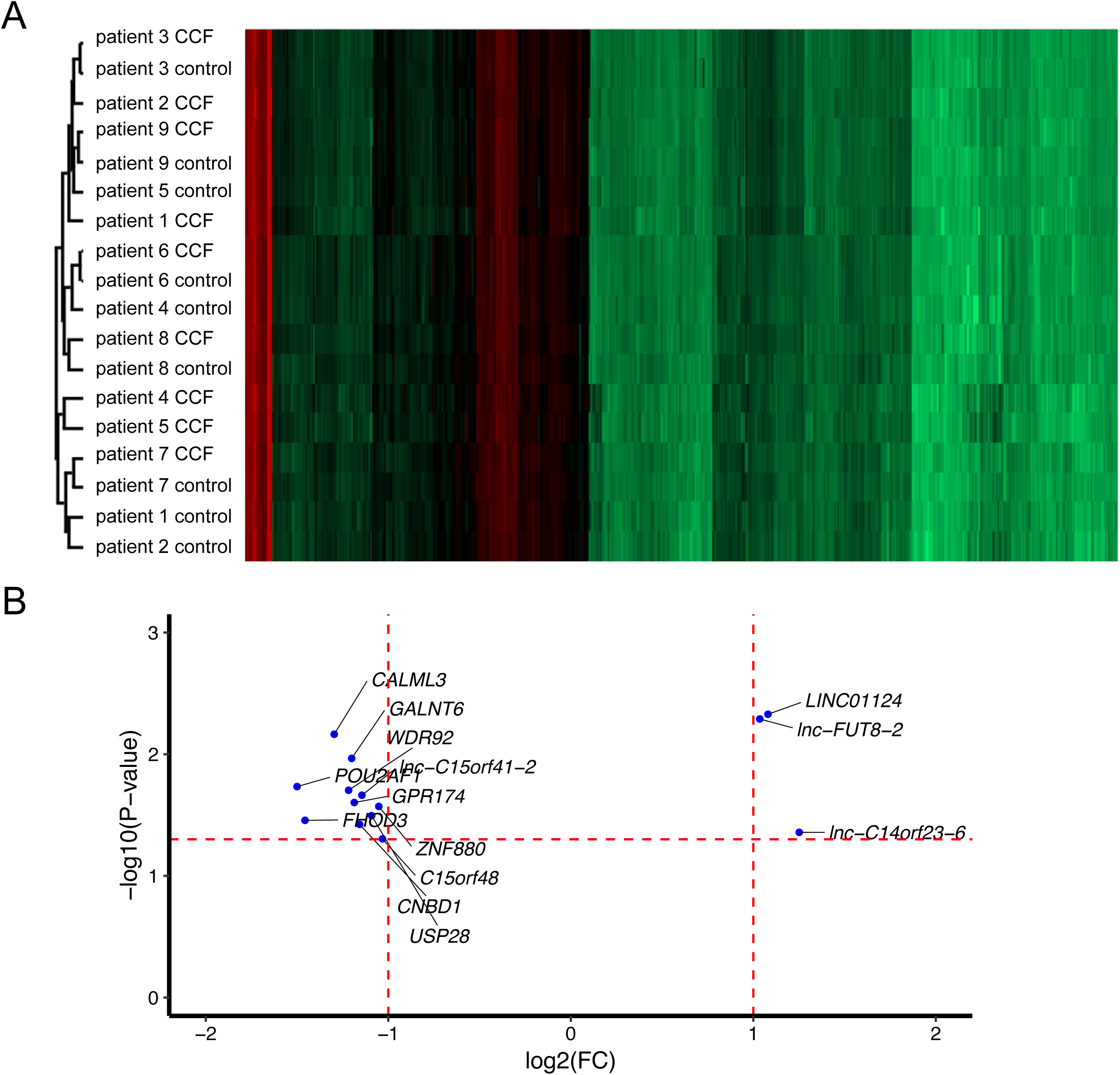
The transcriptome of CCF and control samples from human liver biopsy specimens. Control and CCF samples were laser capture microdissected from cryosections and total RNA was used for microarray analysis. (A) Cluster analysis of the full dataset revealed larger inter-patient differences than differences between CCF and control samples. (B) Scatter plot of transcripts (mRNAs, miRNAs and lncRNAs) with statistically significant differential expression in CCF vs. control samples. More transcripts show lower expression in CCF than in control samples. Horizontal red line: P-value = 0.05; vertical red lines: FC = 2-fold down- or up-regulation, respectively.

Ten of the eleven genes with lower expression in CCF than in control samples code for protein-coding RNAs. The gene with lowest expression was *POU2AF1* (encoding POU class 2 associating factor 1), and is important for the regulation of B-cell maturation [24]. The other down regulated genes have diverse functions in actin binding (FOHD3), calcium-signaling (CALML3), prefoldin-like complex and signaling components (monad/WDR92), protein N-acetylgalactosaminyl transfer (GALNT6), G protein-coupled receptor-mediated signaling (GPR174), nucleotide binding (CNBD1), protein de-ubiquitination (USP28) and as yet unidentified functions (C15orf48 and ZNF880 and the long noncoding RNA lnc-C15orf41-2:1) (Table 1).

### More proteins have reduced expression in CCF compared to control samples

Using mass-spectrometric analysis, we obtained quantitative data of 995 to 2253 proteins in 14 samples (seven control + seven CCF samples from the same patients), with an average of 1474 proteins identified per sample. A bulk of 504 proteins was identified in all 14 samples (Supplementary Figure S1 A). The average number of identified proteins in control and CCF samples did not differ (means/geometric means: 1583/1559 and 1365/1315 proteins; control and CCF, respectively; Supplementary Figure S1 B; student’s t-test, p (unpaired) = 0.300). Also, there was no statistically significant difference between the number of identified proteins from CCF and control samples per patient (Supplementary Figure S1 C; paired student’s t-test, p = 0.137). The normalized log2-transformed expression data approximated normal distribution in all samples (Supplementary Figure S2).

As in the cluster analysis of the microarray data, cluster analysis of the proteomic data could not clearly separate control and CCF samples into distinct groups (Figure 2 A). Therefore, the higher heterogeneity between patients than between CCF and control samples was also present at protein level. Furthermore, more proteins were less expressed in CCF compared to control samples (Figure 2 B and Table 2). The expression of three proteins was decreased more than 2-fold; those were Cytochrome b5 type b (CYB5B, 2.3-fold down), mitochondrial 10 kDa heat shock protein (HSP10/HSPE1; 2.2-fold down) and starch-binding domain-containing protein 1 (STBD1; 2.2-fold down). Further, 19 proteins showed a downregulation of more than 1.5-fold (Figure 2 B and Table 2). In contrast, there were only three proteins with more than 1.5-fold increased expression; these were GSTM4, RAB12 and RAB35. No proteins with >= 2-fold higher expression in CCF than in controls were identified.

**Table 2:** Proteins with statistically significant differential expression in CCF vs control tissue (student’s t-test, N=7)

| Accession | Description (gene symbol) | log2 Ratio<br>(CCF/control) | t-test<br>P-value |
| --- | --- | --- | --- |
| *higher expression in CCF* |  |  |  |
| Q15286 | Ras-related protein Rab-35 (*RAB35*) | 0.9087 | 0.0447 |
| Q6IQ22 | Ras-related protein Rab-12 (*RAB12*) | 0.7768 | 0.0439 |
| Q03013 | Glutathione S-transferase Mu 4 (*GSTM4*) | 0.6028 | 0.0066 |
| *lower expression in CCF* |  |  |  |
| Q12797 | Aspartyl/asparaginyl beta-hydroxylase (*ASPH*) | -0.6097 | 0.0254 |
| Q15008 | 26S proteasome non-ATPase regulatory subunit 6 (*PSMD6*) | -0.6155 | 0.0121 |
| P84098 | 60S ribosomal protein L19 (*RPL19*) | -0.6163 | 0.0086 |
| P61247 | 40S ribosomal protein S3a (*RPS3A*) | -0.6171 | 0.0474 |
| Q13451 | Peptidyl-prolyl cis-trans isomerase FKBP5 (*FKBP*) | -0.6322 | 0.0489 |
| Q9UNF0 | Protein kinase C and casein kinase substrate in neurons protein 2 (*PACSIN2*) | -0.6591 | 0.0327 |
| O95810 | Serum deprivation-response protein (*SDPR*) | -0.6682 | 0.0357 |
| Q16836 | Hydroxyacyl-coenzyme A dehydrogenase, mitochondrial (*HADH*) | -0.6846 | 0.0146 |
| O75821 | Eukaryotic translation initiation factor 3 subunit G (*EIF3G*) | -0.7156 | 0.0333 |
| Q9BVK6 | Transmembrane emp24 domain-containing protein 9 (*TMED9*) | -0.7615 | 0.021 |
| P63208 | S-phase kinase-associated protein 1 (*SKP1*) | -0.7715 | 0.0282 |
| Q15436 | Protein transport protein Sec23A (*SEC23A*) | -0.7936 | 0.0441 |
| Q16629 | Serine/arginine-rich splicing factor 7 (*SRSF7*) | -0.8146 | 0.026 |
| Q95604 | HLA class I histocompatibility antigen, Cw-17 alpha chain (*HLA-C*) | -0.8195 | 0.0429 |
| O75436 | Vacuolar protein sorting-associated protein 26A (*VPS26A*) | -0.878 | 0.0217 |
| Q9NRV9 | Heme-binding protein 1 (*HEBP1*) | -0.9163 | 0.034 |
| O95210 | Starch-binding domain-containing protein 1 (*STBD1*) | -1.1576 | 0.0035 |
| P61604 | 10 kDa heat shock protein, mitochondrial (*HSPE1*) | -1.1642 | 0.0326 |
| O43169 | Cytochrome b5 type B (*CYB5B*) | -1.211 | 0.0157 |

**Figure 2:**
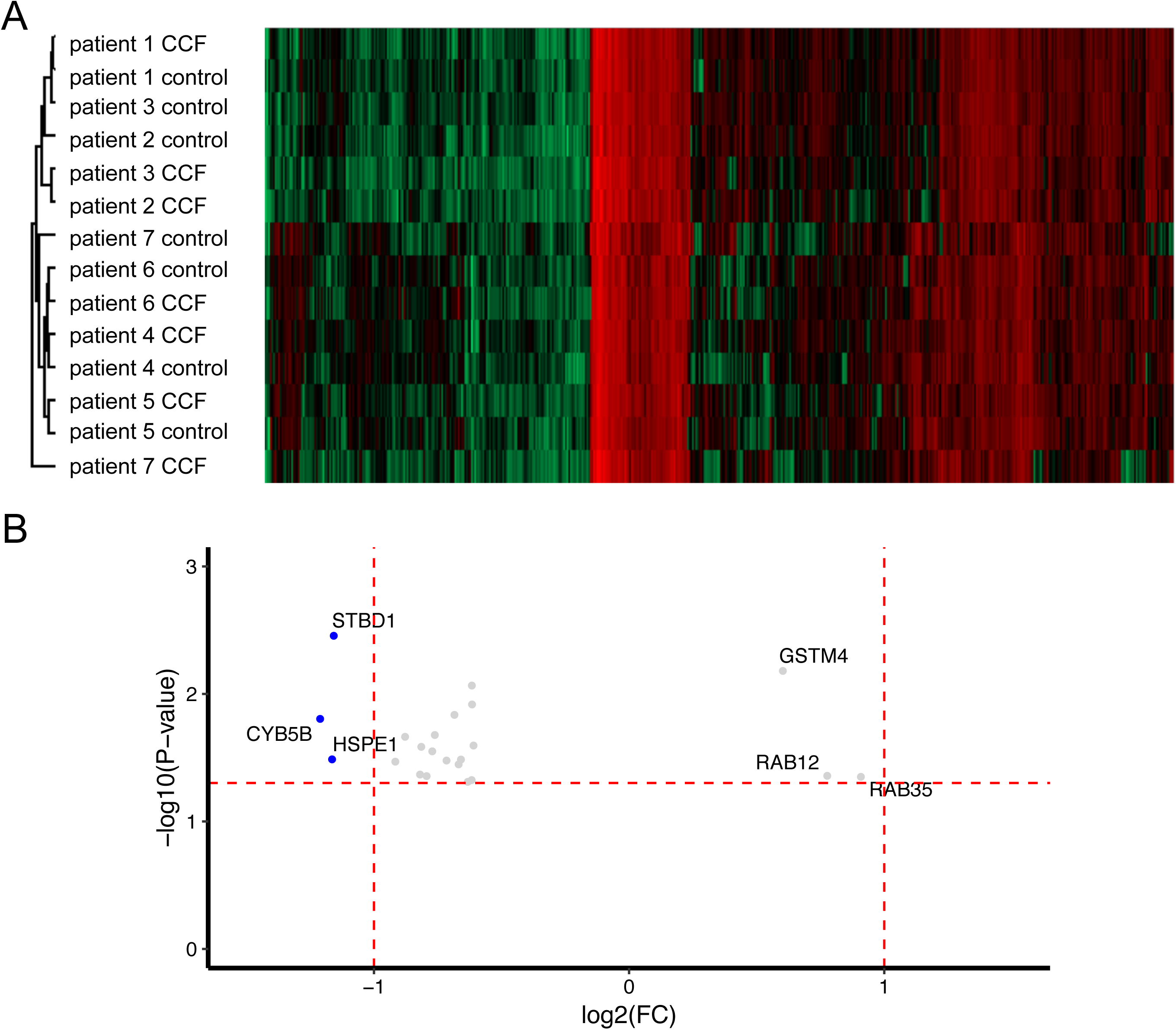
The proteome of CCF and control samples from human liver biopsy specimen. Control and CCF samples were laser capture microdissected from cryosections protein expression was quantified by LC-MS/MS. (A) Cluster analysis of the full dataset revealed larger inter-patient differences than differences between CCF and control samples. (B) Scatter plot of proteins (detection in >= 8 samples) with statistically significant differential expression in CCF vs. control samples. More proteins are less abundant in CCF than in control samples. Horizontal red line: P-value = 0.05; vertical red lines: FC = 2fold down- or upregulation, respectively; proteins significantly different at least 1.5-fold (grey dots) and 2-fold (blue dots) between CCF and control

### Immunohistochemical validation of differential expression of monad/WDR92, USP28, STBD1, CYB5B and HSPE1 in CCF

To validate data obtained with these high-throughput methods, the protein expression of two microarray candidates (monad/WDR92 and USP28) and five proteomics candidates (the downregulated STBD1, CYB5B and HSPE1 as well as the two up-regulated RAB12 and RAB35) was analyzed in liver sections containing CCF using immunohistochemistry. Reduced expression of monad/WDR92, USP28, STBD1, CYB5B and HSPE1 in CCF was confirmed (Figure 3 and Table 3). Increased expression of RAB12 and RAB35 in CCF could not be verified, as in most samples there was no difference between CCF and surrounding tissue or a slight lower expression in CCF (Table 3 and Figure 3). As negative controls we used proteins with unchanged expression like TRAP1, Cullin-3, ACSL4, COPS7A, A-Raf. These proteins had fold changes close to one (1.37, 1.25, 1.11, 1.03 and 1.02, respectively) in the proteomic dataset. Immunohistochemical analysis confirmed no clear difference in expression of these proteins when comparing CCF to surrounding tissue (specimens from 12-15 patients analyzed).

**Table 3:**
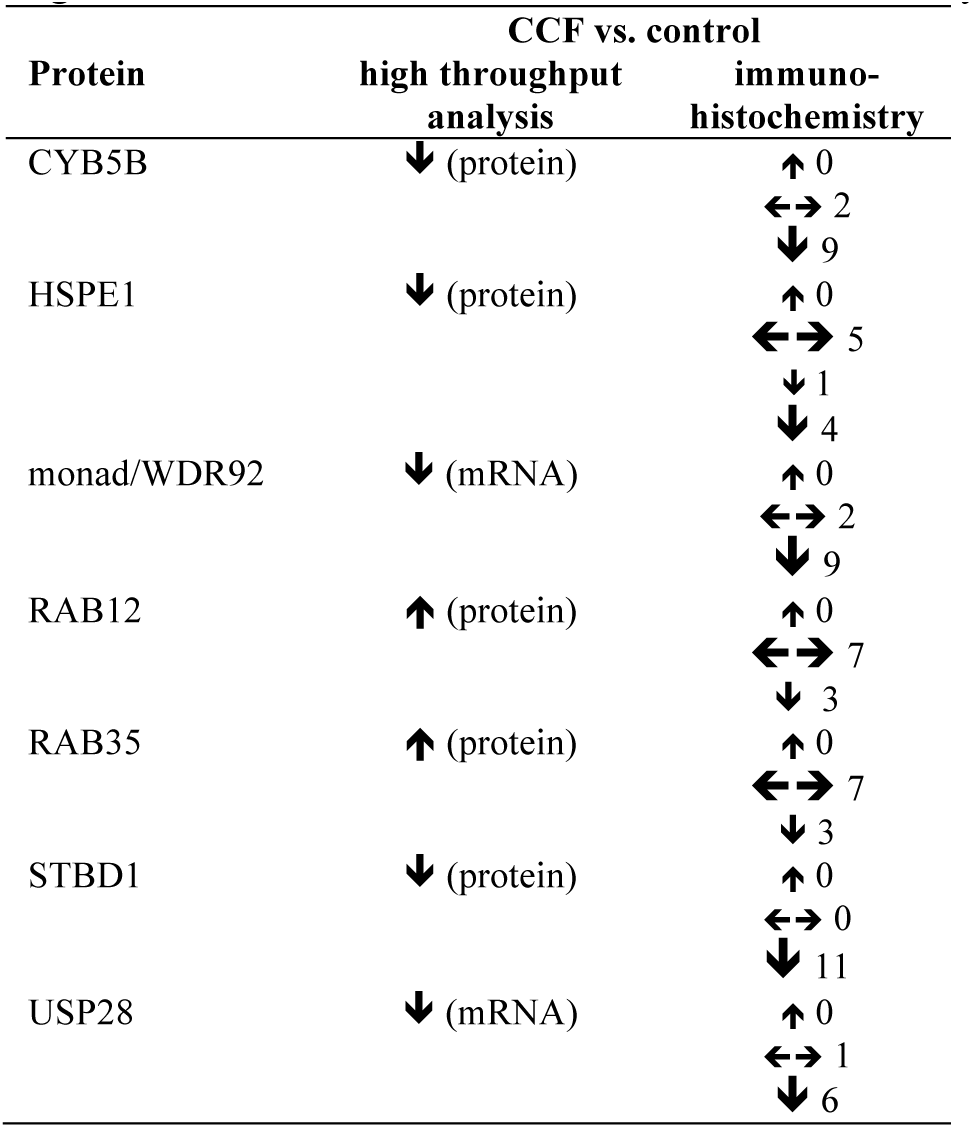
Validation of protein expression as predicted by high-throughput analysis (see Tables 1 and 2 for fold changes) in CCF by immunohistochemical analysis; arrows indicate higher (**↑**), lower (**↓**) and unchanged (**←→**) expression in CCF compared to unaltered liver tissue; brackets around arrows indicate slight differences; numbers at immunohistochemistry column denote the number of individual patient samples analyzed.

| Protein | CCF vs. control |  |
| --- | --- | --- |
|  | high throughput analysis | immuno-histochemistry |
| CYB5B | ↓ (protein) | ↑ 0<br>↔ 2<br>↓ 9 |
| HSPE1 | ↓ (protein) | ↑ 0<br>↔ 5<br>↓ 1<br>↓ 4 |
| monad/WDR92 | ↓ (mRNA) | ↑ 0<br>↔ 2<br>↓ 9 |
| RAB12 | ↑ (protein) | ↑ 0<br>↔ 7<br>↓ 3 |
| RAB35 | ↑ (protein) | ↑ 0<br>↔ 7<br>↓ 3 |
| STBD1 | ↓ (protein) | ↑ 0<br>↔ 0<br>↓ 11 |
| USP28 | ↓ (mRNA) | ↑ 0<br>↔ 1<br>↓ 6 |

**Figure 3:**
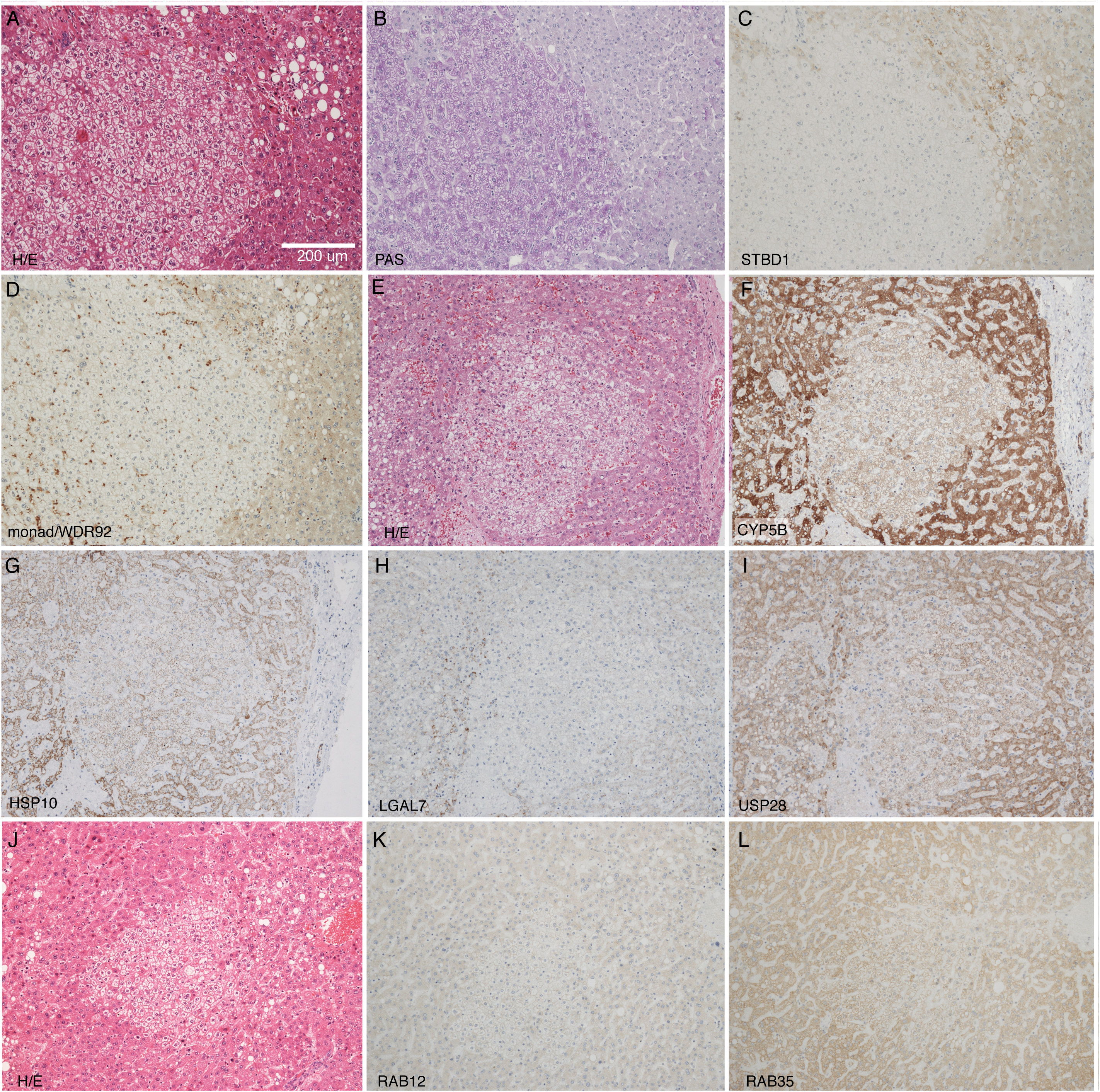
Histochemistry and immunohistochemistry of representative human liver specimens with CCF and neighboring tissue. Hematoxilin/Eosin (H/E) staining (A, E and J), PAS reaction (B), and immunohistochemical detection of STBD1 (C), monad/WDR92 (D), CYP5B (F), HSP10 (G), LGAL7 (H), USP28 (I), RAB12 (K) and RAB35 (L). The scale bar in (A) is applicable to all panels (B-L).

From these results we conclude that the proteomic data from laser capture microdissected samples reflects the protein composition of CCF and surrounding tissue. Our immunohistochemical data of WDR92/monad and USP28 indicate that the observed differences in protein expression in CCF and surrounding tissue could be due to regulation at the transcript levels as the mRNAs coding for these proteins were reduced in CCF according to microarray analysis.

### Loss of STBD1 in mice does not cause glycogen accumulation in the liver

Starch-binding domain-containing protein 1 (STBD1) is N-terminally anchored within the membrane of the endoplasmic reticulum and binds glycogen via its C-terminally located family 20 starch binding module [25]. An Atg8 family interacting motif (AIM) and interaction with GABARAPL1 implicated STBD1 to be involved in autophagic glycogen degradation, so called glycophagy [26]. In a mouse model of Pompe disease, deletion of STBD1 suppressed lysosomal glycogen accumulation, suggesting STDB1 to be involved in transfer of glycogen from the cytoplasm to lysosomes [27].

The reduced expression of STBD1 in CCF could be responsible for the hepatocellular glycogen accumulation. To test whether loss of STBD1 would result in glycogen accumulation in the liver, glycogen concentrations were determined in livers of nine months old male wild type and *Stbd1*-KO mice with access to food throughout or fasted for the last 16 hours. Fasting resulted in significant reductions of glycogen in both, wt and *Stbd1*-KO mice (Figure 4). However, there was no significant difference between wt and *Stbd1*-KO mice at either condition (Figure 4). From these results we conclude that under fasting and normal conditions, loss of STBD1 does not have a significant effect on glycogen degradation in otherwise healthy mice.

**Figure 4:**
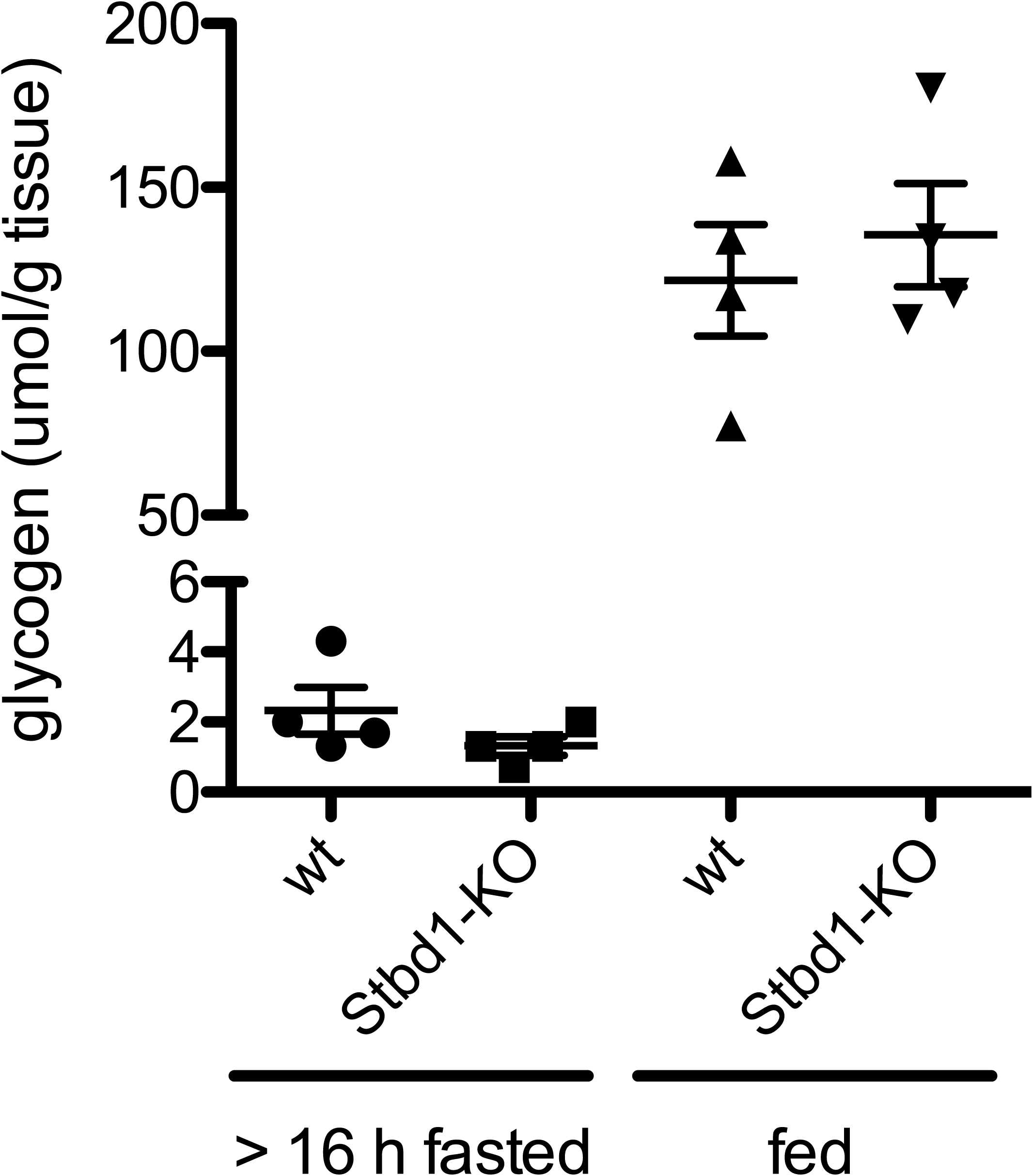
Glycogen concentrations in livers of *Stbd1*-KO mice after more than 16 hours of fasting or in fed state. Male mice (age of 9 months) had access to food ad libitum (fed); the fasted group did not have access to food during 16 hours prior to sampling. N = 4 per group; student’s t-test not significant.

### Diethylnitrosamine (DEN)-induced hepatocellular carcinomas in *Usp28*-KO mice do not accumulate more glycogen than in control mice

Knockout of *Usp28* in mice promotes liver carcinogenesis in diethylnitrosamine (DEN)- injected mice. Although the mechanism was supposed to involve p53 through 53BP1 [28-30] no major impact for this regulatory axis was found in the DEN-induced HCC’s of *Usp28*-KO mice [31]. As USP28 levels were downregulated in the CCF it could mediate its effects at least partially via glycogen regulation. To investigate whether loss of USP28 may affect glycogen metabolism in carcinoma, we compared glycogen accumulation in DEN-induced HCC of wt and *Usp28*-KO mice using PAS staining. However, no difference in PAS staining could be detected suggesting that lack of USP28 does not cause glycogen differences in CCF (Figure 5).

**Figure 5:**
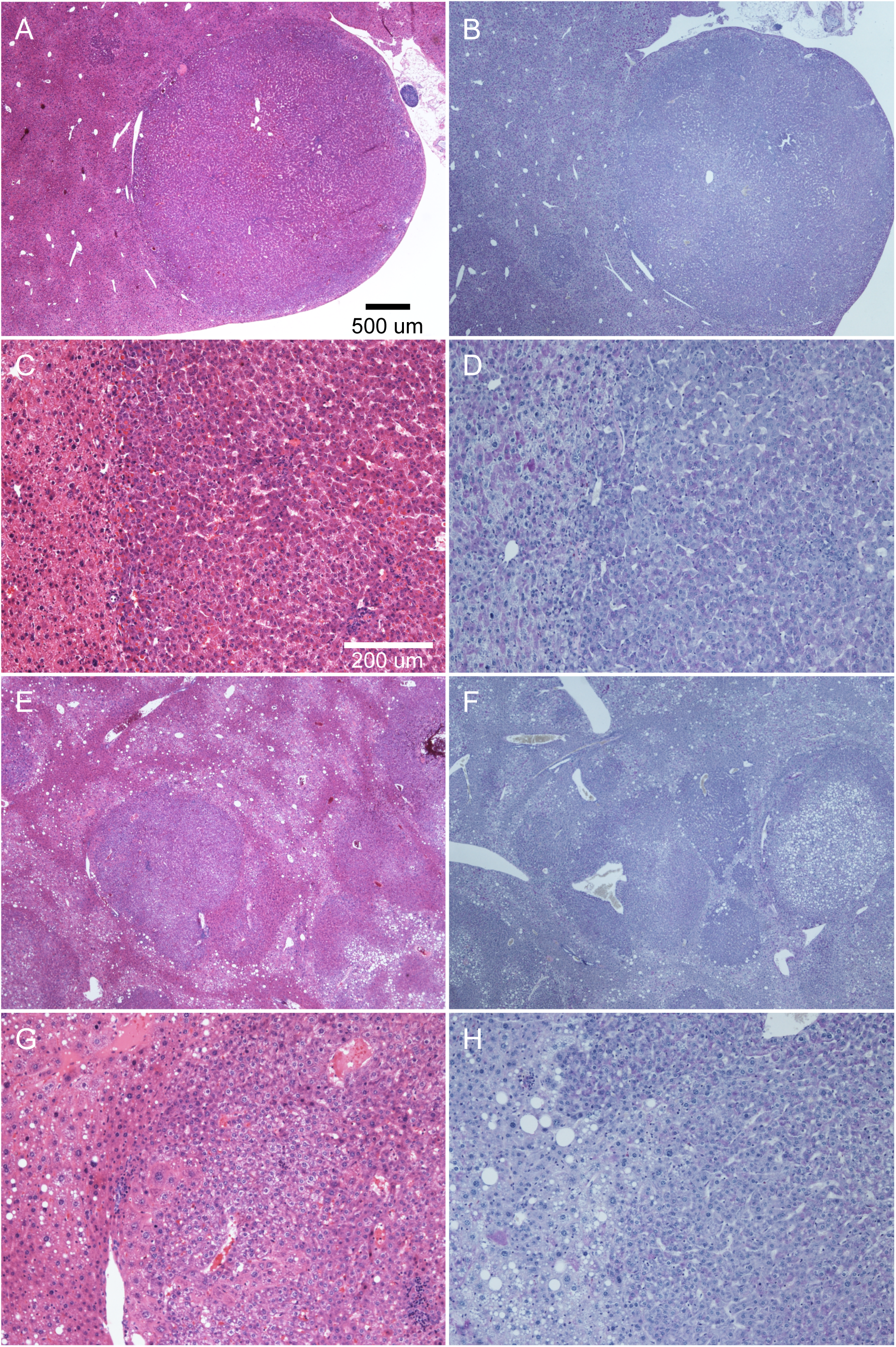
Histochemistry of diethylnitrosamine (DEN)-induced hepatocellular carcinomas of wild type (A-D) and *Usp28* knock out (E-H) mice. H/E stain (A, C, E and G) and PAS reaction (B, D, F and H). Representative images are shown. A, B and E, F are overviews of C, D and G, H respectively; the scale bars in A and C are applicable to B, E and F and D, G and H respectively.

## DISCUSSION

In this study, we identified novel genes and proteins that are differentially regulated in human CCF in comparison to the surrounding liver tissue to better understand the processes leading to glycogen accumulation and the supposed tumor development. We found only few genes/proteins with significant changes higher than two-fold. However, we were able to validate the differential expression of several candidates by immunohistochemistry. Comparing the microarray and the proteomic data, no protein was found for which the respective RNA was changed accordingly. Therefore, these proteins may be not regulated at the transcript level to the same degree-they are regulated at the protein level. It is well known that the level of correlation between RNA expression and protein expression at the genomic level is estimated to be 40-50% [32], and the identified proteins may belong to the 50-60% of proteins that are mainly regulated at the post-transcriptional level.

Notably, in both, the RNA and protein dataset, more genes/proteins were down-regulated than up-regulated in CCF, and the ratios of up-regulated versus down-regulated hits were similar (RNA: 3 up, 11 down (FC cutoff 2fold); protein: 6 up, 22 down (FC cutoff 1.5fold)). This could indicate a general non-specific replacement of cellular components by glycogen. However, we did not find indications that signal intensities of RNA and proteins were generally lower in CCF samples compared with control samples. Hence, we conclude that the differences in RNA and protein levels we observed between CCF and surrounding tissue were due to regulatory processes within cells.

### Non-coding RNAs

We found several non-coding RNAs to be differentially expressed in CCFs compared to controls. Unfortunately, no function is known for any of these. However, lnc-FOXG1-6:17 is an antisense lncRNA of *PRKD1* encoded by two exons (LNCIPEDIA v 5.2, www.lncipedia.org). According to current understanding, antisense transcripts are implicated in gene regulation, which can act in *cis* through transcriptional interference, double-stranded RNA/RNA masking, double-stranded RNA/RNA adenin to inosine editing or double-stranded RNA/RNA interference. In *cis* and *trans* antisense lncRNAs may affect gene expression more globally through chromatin modifications (reviewed in [33]). Hence, it is not clear if and how lnc-FOXG1-6:17 would affect the expression of *PRKD1*, which codes for Serine/threonine-protein kinase D1 (PRKD1). PRKD1 itself is implicated in cell proliferation, cell motility, invasion, protein transport and apoptosis [34] and would represent an interesting candidate in regard to tumor development and progression in CCF. Its RNA and protein expression, though, is low in liver when compared to other tissues (human protein atlas, [35-37]) and its mRNA level was not altered according to our microarray analysis. Hence, a direct and strong effect of lnc-FOXG1-6:17 on *PRKD1* expression seems unlikely.

### STBD1

Glycogen accumulation in CCF is usually explained by decreased gluconeogenesis due to lower activity of the glucose-6-phosphatase increased glycogen synthesis and reduced glycogen degradation mediated by insulin/AKT-signaling [7]. In our current work we observed reduced expression of starch-binding domain-containing protein 1 (STBD1) in CCF compared to control/surrounding tissue in the proteomic dataset and validated this finding by immunohistochemistry. STBD1 is a N-terminally membrane anchored [38] glycogen-binding [25,39] protein that localizes glycogen to perinuclear sites/ER, late endosomes and lysosomes and is most abundantly expressed in muscle and liver [25]. The lower expression of STBD1 we observed in CCF together with its involvement in the lysosomal glycogen degradation pathway [27] suggests that this reduction of STBD1-expression could be an additional factor contributing to glycogen accumulation in CCF. However, a significant glycogen accumulation in livers of *Stbd1*-KO mice was not observed. This was unexpected, as 10% of glycogen degraded in the liver passes through the lysosomal pathway [40] and the loss of STBD1 should result in an appreciable accumulation of glycogen in the liver. One explanation for the lack of this glycogen accumulation is the compensation of STBD1-mediated glycogen translocation into lysosomes by cytoplasmic glycogen degradation Alternative mechanisms of glycogen-translocation are also possible, as the loss of STBD1 in a mouse model of lysosomal glycogen overload (acid alpha-glucosidase-KO mice) did not completely antagonized lysosomal glycogen accumulation [27]. Hence, we conclude that reduction of STBD1 in CCF may only contribute to glycogen accumulation in combination with yet unidentified factors that favor lysosomal glycogen degradation. As large glycogen granulae are preferentially degraded via the lysosomal pathway [40] it would be of interest to determine whether glycogen granulae in CCF are of larger size than in normal liver tissue. An alternative interpretation of the reduction of STBD1 in CCF could be a rerouting of carbohydrate use. Reducing the utilisation of glycogen for lysosomal degradation and glucose export would result in increased intracellular glucose availability. Indeed, cancer cells have been shown to route glucose through glycogen and this glucose was able to support the pentose phosphate pathway more optimally [41,42].

### USP28

In regard to hepatocellular carcinogenesis, USP28 is of particular interest. Functionally, USP28 is a deubiquitination enzyme, catalyzing the deubiquitination of target proteins [43,44], thereby counteracting ubiquitin-dependent proteasomal protein-degradation. Known target proteins of USP28 are 53BP1 [28-30] and claspin [44], MYC [45], LSD1 [46] histone H2A [47] and HIF-1alpha[48]. In particular, lack of USP28 results in an earlier onset and greater tumor burden in a mouse model of chemically (diethylnitrosamine (DEN)) induced hepatocellular carcinoma [31]. The same study also reports that USP28 expression is reduced in patients with hepatocellular carcinoma when comparing carcinoma with control liver tissue of the same patient [31] in line with the data from the current study. Functions of USP28 in different cancer cell types reveal a variety of cancer-relevant effects of USP28: it elicits stem-cell-like characteristics through LSD1-stabilization [46], it acts on p53 and GATA4 affecting cellular senescence [49], it impacts on cell proliferation through deubiquination of histone H2A [47], and it sensitizes cells to DNA-damage via its interactions with 53BP1 and claspin [44]. Finally, USP28 has been targeted in several studies for the treatment of different cancers, such as non-small cell lung cancer, breast cancer, intestinal cancers, gliomas and bladder cancer [50]. Our data (reduced *USP28* mRNA in CCF, validated at protein level by immunohistochemistry) now reveal changes in USP28 expression already in small human hepatocellular foci, supporting the assumption of CCF as very early lesions in hepatocellular carcinogenesis or pre-neoplastic lesions, respectively. However, we did not find any evidence that lack of USP28 would induce glycogen overload in DEN-induced hepatocellular carcinoma. Hence, we assume that reduced expression of USP28 would rather affect the reported p53/senescence pathway axis to promote tumorigenesis and cancerogenesis in clear cell foci. To our knowledge, the alteration of USP28 expression in early clear cell lesions is the first of its kind and it will be interesting to investigate mechansims that trigger the reduced USP28 expression in CCF. Our results of reduced USP28 mRNA expression (microarrays) and reduced USP28 protein expression (IHC)) suggest that the differential regulation is mediated at the transcript level.

## MATERIALS AND METHODS

### Human liver specimens

Liver samples originated from a former cohort [20] without signs of liver cirrhosis obtained from human liver resections taken during surgery. Specimens were from patients (age ranging from 42 to 77 years) with liver metastases of tumors of different origin (two colon carcinomas, two neuroendocrine tumours, one gastrointestinal stromal tumor) or with cholangiocellular carcinoma (n=4)). Tissue samples (1.5 cm x 1.5 cm x 0.5 cm) were collected and frozen in liquid nitrogen-cooled isopentane and stored at −80°C until cryosectioning. Experiments were reviewed and permitted by the ethical committee of the Universitaetsmedizin Greifswald (No. BB 67/10).

### Cryosectioning

Cryosections were made at −20°C to −16°C using a Cryostar NX (Thermo Scientific, Waltham, MA, USA) disinfected with Leica Cryofect Desinfectant Spray (Leica, Wetzlar) before use. All equipment was cleaned with RNase AWAY (Molecular Bio Products, San Diego, USA) and glassware was additionally rinsed with RNase-free water (Aqua B. Braun, Ecotainer, B. Braun Melsungen AG, Melsungen, Germany) and a new blade was used for every sample to reduce risks of cross-contamination. Samples were attached to an object-holder with 0.9% sodium chloride solution (NaCl 0.9%; B. Braun Melsungen AG, Melsungen, Germany). Specimens with clear cell foci were identified by H&E staining of 8 µm sections and validated by PAS reaction. In regions with CCF two to four consecutive cryosections with a thickness of 12 µm were put on membrane-slides (Leica Frame Slides, Nuclease and human nucleic acid free, PET-Membrane, 1,4 µm, Leica, Wetzlar, Germany) and incubated for one minute in 70% ethanol at −16°C. Membrane-slides were collected in a box, vacuum-sealed (Severin FS 3602, Severin Elektrogeräte GmbH, Sundern, Germany) and stored at −80°C until laser microdissection.

### Laser microdissection

CCF and control samples were isolated by laser microdissection using a Leica LMD 6500 System (Leica Microsystems, Wetzlar, Germany) wiped with RNase AWAY (Molecular Bio Products, San Diego, USA). Sample boxes were thawed 30 minutes on ice before staining the cryosections according to a modified H&E-staining protocol. Briefly: sections were incubated in DEPC-water (0.1% diethyl-pyrocarbonate, Sigma-Aldrich, St. Louis, MO, USA) for ten seconds, stained with hemalaun for 50 seconds, and washed for 10 seconds with DEPC-water before staining with eosin for ten seconds. Slides were then incubated in 90% ethanol for 30 seconds.

Material of the same patient and same sample type was collected in one tube and stored on dry ice during collection. For storage, 800µl TRIzol Reagent (Life Technologies, Carlsbad, CA, USA) was added and samples were stored at −80°C.

For microarray analysis nine (N=9) and for proteomic analysis seven sample pairs (N=7) were appropriate.

### RNA and protein isolation

Samples (ca. 90-100 dissected tissue pieces per patient and sample type) were homogenized in liquid nitrogen pre-cooled 4 ml PTFE-vials with one 8 mm stainless steel bead using a Micro-Dismembranator (Sartorius AG, Göttingen) at 2600 RPM for two minutes. The homogenate was transferred into a 1.7 ml centrifuge tube (Sorenson Bioscience Inc., Murray, UT, USA) and the homogenization vials were flushed with another 200 µL Trizol. Samples stored on dry ice were thawed at room temperature for ten minutes, centrifuged at 4 °C and 12000 x g for ten minutes (Heraeus Fresco 17 Centrifuge Refrigerated, Thermo Scientific, Waltham MA, USA) to remove crude debris.

RNA and proteins were isolated using Trizol extraction according to the manufacturers protocol with the following changes. RNA was precipitated with isopropanol over night at −20 °C, RNA was washed twice with 70% ethanol and pellets were resuspended by incubation on water ice for three hours before 30 minutes incubation at room temperature.

Protein pellets were solubilized according to the manufacturers protocols, for smaller pellets 50 µl 1% SDS (Sodium Dodecyl Sulfate, Bio-Rad Laboratories Inc., Hercules, CA, USA) was used.

RNA was quantified using a Nanodrop 8000 (Thermo Scientific, Waltham, MA, USA) and RNA quality was assessed using a Bioanalyzer (Agilent 2100 Bioanalyzer, Agilent Technologies, Santa Clara, CA, USA) (Supplementary Table S1).

### Gel electrophoresis and silver staining

Protein concentrations were determined by Bradford assay (Quick Start Bradford Protein Assay, Bio-Rad Laboratories Inc., Hercules, CA, USA) (Table S1). To assess protein integrity, 13.7-20 µg of total protein per sample was resolved under reducing conditions on Novex NuPAGE gels (4-12% bis-tris protein gels) using LDS sample buffer and MES running buffer according to the manufacturers protocol (Life Technologies, Carlsbad, USA). Gels were stained using the Silver Staining Plus Kit and documented on the ChemiDoc XRS+ system (Bio-Rad Laboratories Inc., Hercules, USA) according to the manufacturers protocol.

### Proteomics-sample preparation and LC-MS/MS

Sample preparation and mass-spectrometric analysis was carried out by the proteomic facility of Porto Conte Ricerche (Alghero, Italy). Protein extracts were subjected to on-filter reduction, alkylation, and trypsin digestion according to the filter-aided sample preparation (FASP) protocol with slight modifications. Briefly, protein extracts were diluted in 8 M urea in Tris HCl 100mM pH 8.8, and buffer was exchanged using Microcon Ultracel YM-10 filtration devices (Millipore, Billerica, MA, USA). Proteins were reduced in 10 mM DTT for 30 min, alkylated in 50 mM iodoacetamide for 20 min, washed five times (3x in 8M urea and 2x in ammonium bicarbonate), before trypsin digestion on the filter (1:100 enzyme-to-protein ratio) at 37 °C overnight. Peptides were collected by centrifugation followed by an additional wash with an elution solution (70% acetonitrile plus 1% formic acid). Finally, the peptide mixture was dried, reconstituted in 0.2% formic acid to a approximate final concentration of 1 µg/µl. Peptide mixture concentration was estimated by measuring absorbance at 280 nm with a NanoDrop 2000 spectrophotometer (Thermo Scientific, San Jose, CA, USA) and a standard curve made from MassPREP *E. Coli* Digest Standard (Waters, Milford, MA, USA).

LC-MS/MS analyses were carried out using a Q Exactive mass spectrometer (Thermo Scientific) interfaced with an UltiMate 3000 RSLCnano LC system (Thermo Scientific). After loading, peptide mixtures (4 *µ*g per run) were concentrated and desalted on a trapping pre-column (Acclaim PepMap C18, 75 *µ*m × 2 cm nanoViper, 3 *µ*m, 100 Å, Thermo Scientific), using 0.2% formic acid at a flow rate of 5 *µ*l/min. The peptide separation was performed at 35 °C using a C18 column (EASY-Spray column, 15cm x 75 *µ*m ID, PepMap C18, 3*µ*m, Thermo Scientific) at a flow rate of 300 nL/min, using a 485 min gradient from 1 to 50% eluent B (0.2% formic acid in 95% acetonitrile) in eluent A (0.2% formic acid in 5% acetonitrile). MS data were acquired using a data-dependent top10 method dynamically choosing the most abundant precursor ions from the survey scan, under direct control of the Xcalibur software (version 1.0.2.65 SP2), where a full-scan spectrum (from 300 to 1,700 m/z) was followed by tandem mass spectra (MS/MS). The instrument was operated in positive mode with a spray voltage of 1.8 kV and a capillary temperature of 275°C. Survey and MS/MS scans were performed in the Orbitrap with resolution of 70,000 and 17,500 at 200 m/z, respectively. The automatic gain control was set to 1,000,000 ions and the lock mass option was enabled on a protonated polydimethylcyclosiloxane background ion as internal recalibration for accurate mass measurements. The dynamic exclusion was set to 30 seconds. Higher Energy Collisional Dissociation (HCD), performed at the far side of the C-trap, was used as fragmentation method, by applying a 25 eV value for normalized collision energy, an isolation width of m/z 2.0. Nitrogen was used as the collision gas.

Peptide identification was performed using Proteome Discoverer (version 1.4; Thermo Scientific) using Sequest-HT as search engine for protein identification, according to the following criteria: Database UniprotKB, taxonomy human (release 2014_10); Precursor mass tolerance: 10 ppm; Fragment mass tolerance: 0.02 Da; Static modification: cysteine carbamidomethylation; Dynamic modification: methionine oxidation, and Percolator for peptide validation (FDR < 1% based on peptide q-value). Results were filtered in order to keep only rank 1 peptides, and protein grouping was allowed according to the maximum parsimony principle.

Protein abundance was expressed by means of the normalized spectral abundance factor (NSAF). NSAF was calculated as follows: NSAF=SAFi/N, where subscript i denotes a protein identity and N is the total number of proteins, while SAF is a protein spectral abundance factor that is defined as the protein spectral counts divided by its length (number of residues or molecular weight). In this approach, the spectral counts of each protein were divided by its length and normalized to the average number of spectral counts in a given analysis. In order to eliminate discontinuity due to SpC=0, a correction factor, set to 2, was used. The NSAF log ratio (RNSAF) was calculated according to the following formula: RNSAF= log2(NSAF1/NSAF2) where RNSAF is the log2 ratio of the abundance of a protein in sample groups 1 (Preneo, NSAF1) and 2 (Norm, NSAF2).

Proteins showing RNSAF > 0.5 or < −0.5 were considered as differentially abundant between groups. A two-tailed t-test was applied, using in house software, in order to evaluate the statistical significance of differences between groups.

### Microarray analysis

Processing of purified RNA for microarray analysis and microarray analysis was carried out at OakLabs (Hennigsdorf, Germany) according to their standard procedures. Briefly. RNA concentrations were between 17 and 66 ng/uL in volumes of 30 or 40 uL H_2_O with integrity numbers (RIN, Bioanalyzer, Agilent Technologies, USA) between 6.3 and 7.9 (**Table S1**). RNA was labeled using the Low Input QuickAmp Labeling Kit (Agilent Technologies, USA) and cRNA was hybridized with ArrayXS Human (Oaklabs, Germany) at 65 °C for 17 hours using the Agilent Gene Expression Hybridization Kit (Agilent Technologies, USA), washed once with Agilent Gene Expression Wash Buffer 1 for one minute at room temperature, followed by a second wash with preheated (37 °C) Gene Expression Wash Buffer 2 for one minute.

Microarrays were scanned with a SureScan Microarray Scanner (Agilent Technologies, USA) and Agilent’s Feature Extraction software was used to detect features. Signals from control probes were removed and means of signals from replicate probes and of signals from all probes of a target were determined before normalization of the background subtracted signals. Data of all samples was quantile normalized using ranked mean quartiles [51]. Normalized data was statistically analyzed by paired ANOVA (clear cell foci vs control samples from the same patient).

### Cluster analysis and plotting of omics-data

Cluster analysis of proteomic and microarray data was done using Perseus Software (version 1.6.0.7,). Micro array data: quartile normed data was filtered by intensity (signal >=10, in 8 out of 9 samples within one group; groups: normal and CCF). Settings for hierarchical clustering: Distance - Euclidean; Linkage - Average (process with k-means; number of clusters 300; maximal number of iterations 10; number of restarts 1). Data plotting was done using R (3.5.2) with the following packages: ggplot2, ggrepel and gridExtra.

### Immunohistochemistry and histochemistry

Histochemistry of formaldehyde fixed and paraffin embedded specimen was performed as previously described [14].

Immunohistochemistry was carried out on formaldehyde fixed and paraffin embedded specimen according to standard immunohistochemical protocols for de-paraffination and embedding. Endogenous peroxidase was blocked with Novocastra™Peroxidase Block (#RE7101, Leica Biosystems) for 15 minutes at room temperature and sections were blocked with Universal Block (Dako) for 20 minutes. Primary antibodies and unmasking techniques are listed in Supplementary Table S2. LSAB2 System-HRP (#K0675, Dako) and Liquid DAB+ Substrate Chromogen System (#K3468, Dako) were used for signal amplification and staining.

### Mouse models

*Stbd1*-KO mice [27] and the respective control animals were bred and housed at Duke University, USA. All animal procedures were done in accordance with Duke University Institutional Animal Care and Use Committee-approved guidelines.

Sections of *Usp28*-KO mice were prepared as described [31].

### Glycogen quantification in mouse liver

Glycogen was quantified in mouse liver as described previously [52].

## AUTHOR CONTRIBUTIONS

Conception or design of the work: S.R., D.F.C. and F.D.

Data collection: K.W., J.R., C.M., D.M., and A.C.

Data analysis and interpretation: S.R., F.D., D.F.C. and C.M.

Drafting the article: S.R. and C.M.

Critical revision of the article: S.R., F.D., T.K., D.F.C. and C.M.

Final approval of the version to be published: S.R., F.D., D.F.C., A.C., J.R., T.K., B.S., D.M., K.W., and C.M.

## ACKNOWLEDGMENTS

The authors thank Rebekka Icke and Anke Wolter for technical assistance.

## REFERENCES

1. Franco LM, Krishnamurthy V, Bali D, Weinstein DA, Arn P, Clary B, Boney A, Sullivan J, Frush DP, Chen Y-T, Kishnani PS. Hepatocellular carcinoma in glycogen storage disease type Ia: a case series. J Inherit Metab Dis. John Wiley & Sons, Ltd; 2005; 28: 153–62. doi: 10.1007/s10545-005-7500-2.

2. Marengo A, Rosso C, Bugianesi E. Liver Cancer: Connections with Obesity, Fatty Liver, and Cirrhosis. Annu Rev Med. Annual Reviews; 2016; 67: 103–17. doi: 10.1146/annurev-med-090514-013832.

3. Davila JA, Morgan RO, Shaib Y, McGlynn KA, El-Serag HB. Diabetes increases the risk of hepatocellular carcinoma in the United States: a population based case control study. Gut. BMJ Publishing Group; 2005; 54: 533–9. doi: 10.1136/gut.2004.052167.

4. Vigneri R, Goldfine ID, Frittitta L. Insulin, insulin receptors, and cancer. J Endocrinol Invest. Springer International Publishing; 2016; 39: 1365–76. doi: 10.1007/s40618-016-0508-7.

5. Libbrecht L, Desmet V, Roskams T. Preneoplastic lesions in human hepatocarcinogenesis. Liver Int. John Wiley & Sons, Ltd (10.1111); 2005; 25: 16–27. doi: 10.1111/j.1478-3231.2005.01016.x.

6. Evert M, Dombrowski F. [Hepatocellular carcinoma in the non-cirrhotic liver]. Pathologe. 2008; 29: 47–52. doi: 10.1007/s00292-007-0953-3.

7. Bannasch P, Ribback S, Su Q, Mayer D. Clear cell hepatocellular carcinoma: origin, metabolic traits and fate of glycogenotic clear and ground glass cells. HBPD INT. 2017; 16: 570–94. doi: 10.1016/S1499-3872(17)60071-7.

8. Bannasch P. Pathogenesis of hepatocellular carcinoma: sequential cellular, molecular, and metabolic changes. Prog Liver Dis. 1996; 14: 161–97.

9. Bannasch P, Mayer D, Hacker HJ. Hepatocellular glycogenosis and hepatocarcinogenesis. Biochim Biophys Acta. 1980; 605: 217–45.

10. Williams GM. The pathogenesis of rat liver cancer caused by chemical carcinogens. Biochim Biophys Acta. 1980; 605: 167–89.

11. R Pitot HC. Altered hepatic foci: their role in murine hepatocarcinogenesis. Annu Rev Pharmacol Toxicol. Annual Reviews 4139 El Camino Way, P.O. Box 10139, Palo Alto, CA 94303-0139, USA; 1990; 30: 465–500. doi: 10.1146/annurev.pa.30.040190.002341.

12. Dombrowski F, Filsinger E, Bannasch P, Pfeifer U. Altered liver acini induced in diabetic rats by portal vein islet isografts resemble preneoplastic hepatic foci in their enzymic pattern. Am J Pathol. American Society for Investigative Pathology; 1996; 148: 1249–56.

13. Dombrowski F, Mathieu C, Evert M. Hepatocellular neoplasms induced by low-number pancreatic islet transplants in autoimmune diabetic BB/Pfd rats. Cancer Res. American Association for Cancer Research; 2006; 66: 1833–43. doi: 10.1158/0008-5472.CAN-05-2787.

14. Dombrowski F, Bannasch P, Pfeifer U. Hepatocellular neoplasms induced by low-number pancreatic islet transplants in streptozotocin diabetic rats. Am J Pathol. American Society for Investigative Pathology; 1997; 150: 1071–87.

15. Dombrowski F, Jost CM, Manekeller S, Evert M. Cocarcinogenic effects of islet hormones and N-nitrosomorpholine in hepatocarcinogenesis after intrahepatic transplantation of pancreatic islets in streptozotocin-diabetic rats. Cancer Res. American Association for Cancer Research; 2005; 65: 7013–22. doi: 10.1158/0008-5472.CAN-05-0122.

16. Ribback S, Cigliano A, Kroeger N, Pilo MG, Terracciano L, Burchardt M, Bannasch P, Calvisi DF, Dombrowski F. PI3K/AKT/mTOR pathway plays a major pathogenetic role in glycogen accumulation and tumor development in renal distal tubules of rats and men. Oncotarget. 2015; 6: 13036–48. doi: 10.18632/oncotarget.3675.

17. Evert M, Calvisi DF, Evert K, De Murtas V, Gasparetti G, Mattu S, Destefanis G, Ladu S, Zimmermann A, Delogu S, Thiel S, Thiele A, Ribback S, et al. V-AKT murine thymoma viral oncogene homolog/mammalian target of rapamycin activation induces a module of metabolic changes contributing to growth in insulin-induced hepatocarcinogenesis. Hepatology. John Wiley & Sons, Ltd; 2012; 55: 1473–84. doi: 10.1002/hep.25600.

18. Calvisi DF, Wang C, Ho C, Ladu S, Lee SA, Mattu S, Destefanis G, Delogu S, Zimmermann A, Ericsson J, Brozzetti S, Staniscia T, Chen X, et al. Increased lipogenesis, induced by AKT-mTORC1-RPS6 signaling, promotes development of human hepatocellular carcinoma. Gastroenterology. 2011; 140: 1071–83. doi: 10.1053/j.gastro.2010.12.006.

19. Calvisi DF, Ladu S, Gorden A, Farina M, Conner EA, Lee J-S, Factor VM, Thorgeirsson SS. Ubiquitous activation of Ras and Jak/Stat pathways in human HCC. Gastroenterology. 2006; 130: 1117–28. doi: 10.1053/j.gastro.2006.01.006.

20. Ribback S, Calvisi DF, Cigliano A, Sailer V, Peters M, Rausch J, Heidecke C-D, Birth M, Dombrowski F. Molecular and metabolic changes in human liver clear cell foci resemble the alterations occurring in rat hepatocarcinogenesis. J Hepatol. 2013; 58: 1147–56. doi: 10.1016/j.jhep.2013.01.013.

21. Ribback S, Sonke J, Lohr A, Frohme J, Peters K, Holm J, Peters M, Cigliano A, Calvisi DF, Dombrowski F. Hepatocellular glycogenotic foci after combined intraportal pancreatic islet transplantation and knockout of the carbohydrate responsive element binding protein in diabetic mice. Oncotarget. Impact Journals; 2017; 8: 104315–29. doi: 10.18632/oncotarget.22234.

22. Li L, Che L, Tharp KM, Park H-M, Pilo MG, Cao D, Cigliano A, Latte G, Xu Z, Ribback S, Dombrowski F, Evert M, Gores GJ, et al. Differential requirement for de novo lipogenesis in cholangiocarcinoma and hepatocellular carcinoma of mice and humans. Hepatology. John Wiley & Sons, Ltd; 2016; 63: 1900–13. doi: 10.1002/hep.28508.

23. Che L, Pilo MG, Cigliano A, Latte G, Simile MM, Ribback S, Dombrowski F, Evert M, Chen X, Calvisi DF. Oncogene dependent requirement of fatty acid synthase in hepatocellular carcinoma. Cell Cycle. Taylor & Francis; 2017; 16: 499–507. doi: 10.1080/15384101.2017.1282586.

24. Zhao C, Inoue J, Imoto I, Otsuki T, Iida S, Ueda R, Inazawa J. POU2AF1, an amplification target at 11q23, promotes growth of multiple myeloma cells by directly regulating expression of a B-cell maturation factor, TNFRSF17. Oncogene. Nature Publishing Group; 2008; 27: 63–75. doi: 10.1038/sj.onc.1210637.

25. Jiang S, Heller B, Tagliabracci VS, Zhai L, Irimia JM, Depaoli-Roach AA, Wells CD, Skurat AV, Roach PJ. Starch binding domain-containing protein 1/genethonin 1 is a novel participant in glycogen metabolism. J Biol Chem. American Society for Biochemistry and Molecular Biology; 2010; 285: 34960–71. doi: 10.1074/jbc.M110.150839.

26. Jiang S, Wells CD, Roach PJ. Starch-binding domain-containing protein 1 (Stbd1) and glycogen metabolism: Identification of the Atg8 family interacting motif (AIM) in Stbd1 required for interaction with GABARAPL1. Biochem Biophys Res Commun. 2011; 413: 420–5. doi: 10.1016/j.bbrc.2011.08.106.

27. Sun T, Yi H, Yang C, Kishnani PS, Sun B. Starch Binding Domain-containing Protein 1 Plays a Dominant Role in Glycogen Transport to Lysosomes in Liver. J Biol Chem. 2016; 291: 16479–84. doi: 10.1074/jbc.C116.741397.

28. Lambrus BG, Daggubati V, Uetake Y, Scott PM, Clutario KM, Sluder G, Holland AJ. A USP28-53BP1-p53-p21 signaling axis arrests growth after centrosome loss or prolonged mitosis. J Cell Biol. 2016; 214: 143–53. doi: 10.1083/jcb.201604054.

29. Fong CS, Mazo G, Das T, Goodman J, Kim M, O’Rourke BP, Izquierdo D, Tsou M-FB. 53BP1 and USP28 mediate p53-dependent cell cycle arrest in response to centrosome loss and prolonged mitosis. Elife. eLife Sciences Publications Limited; 2016; 5: E1491. doi: 10.7554/eLife.16270.

30. Meitinger F, Anzola JV, Kaulich M, Richardson A, Stender JD, Benner C, Glass CK, Dowdy SF, Desai A, Shiau AK, Oegema K. 53BP1 and USP28 mediate p53 activation and G1 arrest after centrosome loss or extended mitotic duration. J Cell Biol. 2016; 214: 155–66. doi: 10.1083/jcb.201604081.

31. Richter K, Paakkola T, Mennerich D, Kubaichuk K, Konzack A, Kippari HA, Kozlova N, Koivunen P, Haapasaari K-M, Jukkola-Vuorinen A, Teppo H-R, Dimova EY, Bloigu R, et al. USP28 Deficiency Promotes Breast and Liver Carcinogenesis as well as Tumor Angiogenesis in a HIF-independent Manner. Mol Cancer Res. American Association for Cancer Research; 2018; 16: 1000–12. doi: 10.1158/1541-7786.MCR-17-0452.

32. Vogel C, Marcotte EM. Insights into the regulation of protein abundance from proteomic and transcriptomic analyses. Nat Rev Genet. Nature Publishing Group; 2012; 13: 227–32. doi: 10.1038/nrg3185.

33. Latgé G, Poulet C, Bours V, Josse C, Jerusalem G. Natural Antisense Transcripts: Molecular Mechanisms and Implications in Breast Cancers. Int J Mol Sci. Multidisciplinary Digital Publishing Institute; 2018; 19: 123. doi: 10.3390/ijms19010123.

34. Youssef I, Ricort J-M. Deciphering the role of protein kinase D1 (PKD1) in cellular proliferation. Mol Cancer Res. 2019; : molcanres.0125.2019. doi: 10.1158/1541-7786.MCR-19-0125.

35. Uhlen M, Zhang C, Lee S, Sjöstedt E, Fagerberg L, Bidkhori G, Benfeitas R, Arif M, Liu Z, Edfors F, Sanli K, Feilitzen von K, Oksvold P, et al. A pathology atlas of the human cancer transcriptome. Science. American Association for the Advancement of Science; 2017; 357: eaan2507. doi: 10.1126/science.aan2507.

36. Thul PJ, Åkesson L, Wiking M, Mahdessian D, Geladaki A, Ait Blal H, Alm T, Asplund A, Björk L, Breckels LM, Bäckström A, Danielsson F, Fagerberg L, et al. A subcellular map of the human proteome. Science. American Association for the Advancement of Science; 2017; 356: eaal3321. doi: 10.1126/science.aal3321.

37. Uhlen M, Fagerberg L, Hallström BM, Lindskog C, Oksvold P, Mardinoglu A, Sivertsson Å, Kampf C, Sjöstedt E, Asplund A, Olsson I, Edlund K, Lundberg E, et al. Proteomics. Tissue-based map of the human proteome. Science. American Association for the Advancement of Science; 2015; 347: 1260419–9. doi: 10.1126/science.1260419.

38. Bouju S, Lignon MF, Piétu G, Le Cunff M, Léger JJ, Auffray C, Dechesne CA. Molecular cloning and functional expression of a novel human gene encoding two 41-43 kDa skeletal muscle internal membrane proteins. Biochem J. 1998; 335 (Pt 3): 549–56. doi: 10.1042/bj3350549.

39. Janeček Š. A motif of a microbial starch-binding domain found in human genethonin. Bioinformatics. 2002; 18: 1534–7. doi: 10.1093/bioinformatics/18.11.1534.

40. Prats C, Graham TE, Shearer J. The dynamic life of the glycogen granule. J Biol Chem. American Society for Biochemistry and Molecular Biology; 2018; 293: 7089–98. doi: 10.1074/jbc.R117.802843.

41. Klimek F, Mayer D, Bannasch P. Biochemical microanalysis of glycogen content and glucose-6-phosphate dehydrogenase activity in focal lesions of the rat liver induced by N-nitrosomorpholine. Carcinogenesis. 1984; 5: 265–8. doi: 10.1093/carcin/5.2.265.

42. Favaro E, Bensaad K, Chong MG, Tennant DA, Ferguson DJP, Snell C, Steers G, Turley H, Li J-L, Günther UL, Buffa FM, McIntyre A, Harris AL. Glucose Utilization via Glycogen Phosphorylase Sustains Proliferation and Prevents Premature Senescence in Cancer Cells. Cell Metabolism. 2012; 16: 751–64. doi: 10.1016/j.cmet.2012.10.017.

43. Valero R, Bayés M, Francisca Sánchez-Font M, González-Angulo O, Gonzàlez-Duarte R, Marfany G. Characterization of alternatively spliced products and tissue-specific isoforms of USP28 and USP25. Genome Biol. BioMed Central; 2001; 2: RESEARCH0043–10. doi: 10.1186/gb-2001-2-10-research0043.

44. Zhang D, Zaugg K, Mak TW, Elledge SJ. A role for the deubiquitinating enzyme USP28 in control of the DNA-damage response. Cell. 2006; 126: 529–42. doi: 10.1016/j.cell.2006.06.039.

45. Popov N, Wanzel M, Madiredjo M, Zhang D, Beijersbergen R, Bernards R, Moll R, Elledge SJ, Eilers M. The ubiquitin-specific protease USP28 is required for MYC stability. Nat Cell Biol. Nature Publishing Group; 2007; 9: 765–74. doi: 10.1038/ncb1601.

46. Wu Y, Wang Y, Yang XH, Kang T, Zhao Y, Wang C, Evers BM, Zhou BP. The deubiquitinase USP28 stabilizes LSD1 and confers stem-cell-like traits to breast cancer cells. Cell Rep. 2013; 5: 224–36. doi: 10.1016/j.celrep.2013.08.030.

47. Li F, Han H, Sun Q, Liu K, Lin N, Xu C, Zhao Z, Zhao W. USP28 regulates deubiquitination of histone H2A and cell proliferation. Exp Cell Res. 2019; 379: 11–8. doi: 10.1016/j.yexcr.2019.03.026.

48. Flügel D, Görlach A, Kietzmann T. GSK-3β regulates cell growth, migration, and angiogenesis via Fbw7 and USP28-dependent degradation of HIF-1α. Blood. 2012; 119: 1292–301. doi: 10.1182/blood-2011-08-375014.

49. Mazzucco AE, Smogorzewska A, Kang C, Luo J, Schlabach MR, Xu Q, Patel R, Elledge SJ. Genetic interrogation of replicative senescence uncovers a dual role for USP28 in coordinating the p53 and GATA4 branches of the senescence program. Genes Dev. Cold Spring Harbor Lab; 2017; 31: 1933–8. doi: 10.1101/gad.304857.117.

50. Wang X, Liu Z, Zhang L, Yang Z, Chen X, Luo J, Zhou Z, Mei X, Yu X, Shao Z, Feng Y, Fu S, Zhang Z, et al. Targeting deubiquitinase USP28 for cancer therapy. Cell Death Dis. Nature Publishing Group; 2018; 9: 186–10. doi: 10.1038/s41419-017-0208-z.

51. Bolstad BM, Irizarry RA, Astrand M, Speed TP. A comparison of normalization methods for high density oligonucleotide array data based on variance and bias. Bioinformatics. 2003; 19: 185–93. doi: 10.1093/bioinformatics/19.2.185.

52. Yi H, Thurberg BL, Curtis S, Austin S, Fyfe J, Koeberl DD, Kishnani PS, Sun B. Characterization of a canine model of glycogen storage disease type IIIa. Dis Model Mech. 2012; 5: 804–11. doi: 10.1242/dmm.009712.

