## Supplementary tables and figures for "Transcriptomic and Proteomic analysis of clear cell foci (CCF) in the human non-cirrhotic liver identifies several differentially expressed genes and proteins with functions in cancer cell biology and glycogen metabolism"

2 - current affilition
Uppsala University, Department of Immunology, Genetics and Pathology, BMC, Husargatan 3 751 24 Uppsala, Sweden

3 - current affiliation
Regensburg Universitaetsklinikum, Institut fuer Pathologie, Franz-Josef-Strauß-Allee 11, 93053 Regensburg, Germany

4 - Division of Medical Genetics, Department of Pediatrics, Duke University Medical Center, Durham, North Carolina 27710, USA

5 - Faculty of Biochemistry and Molecular Medicine, University of Oulu, Oulu, Finland.

6 - Biocenter Oulu, University of Oulu, Oulu, Finland.

Corresponding author:
Silvia Ribback

Universitaetsmedizin Greifswald
Institut fuer Pathologie
Friedrich-Loeffler-Str. 23e
17475 Greifswald, Germany

### SUPPLEMENTARY FIGURES

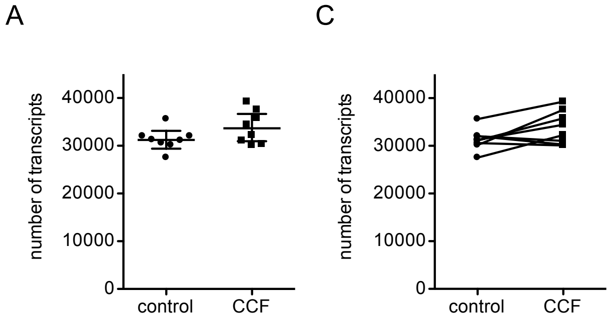


**Supplementary Figure S1:** (**A**) Number of transcripts above threshold per sample and sample-group (control and clear cell foci (CCF)); data of individual samples are shown as dots (controls) and squares (CCF) and geometric means and 95% confidence interval are indicated by lines. (**B**) Number of transcripts expressed above no-expression threshold per sample and sample-group (control and clear cell foci (CCF)); data of individual samples are shown as dots (controls) and squares (CCF) and data of each patient is connected by a line.

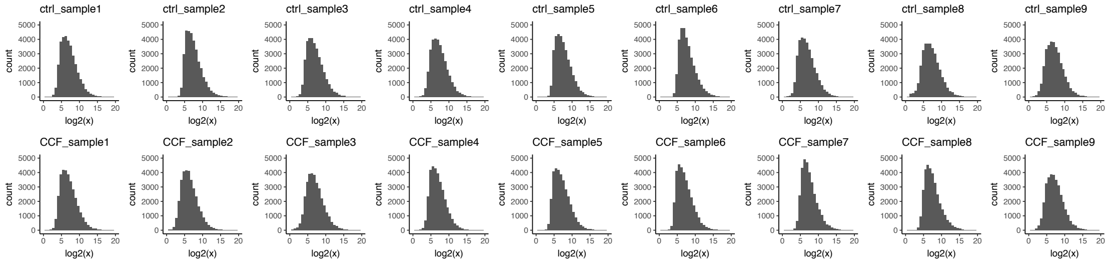


**Supplementary Figure S2:** The distribution of log2 transformed gene expression data approximates normal distribution in all samples and is similar between samples from the same patient. ctrl = control samples, CCF = clear cell foci samples, Sample1-Sample9 denote different patients.

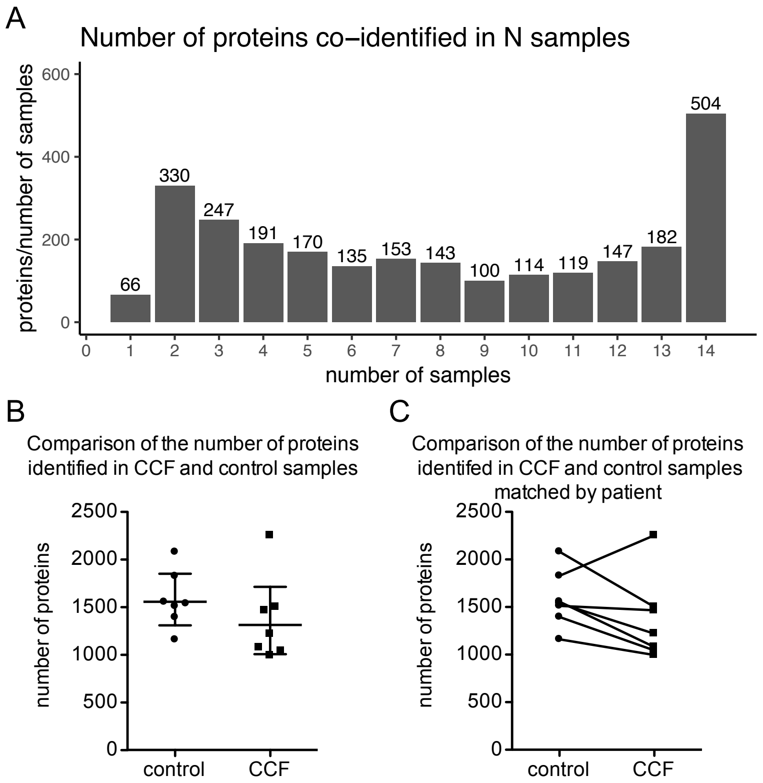


**Supplementary Figure S1:** (**A**) Number of proteins co-identified in the indicated number of samples. (**B**) Number of proteins identified per sample and sample-group (control and clear cell foci (CCF)); data of individual samples are shown as dots (controls) and squares (CCF) and geometric means and 95% confidence interval are indicated by lines. (**C**) Number of proteins identified per sample and sample-group (control and clear cell foci (CCF)); data of individual samples are shown as dots (controls) and squares (CCF) and data of each patient is connected by a line.

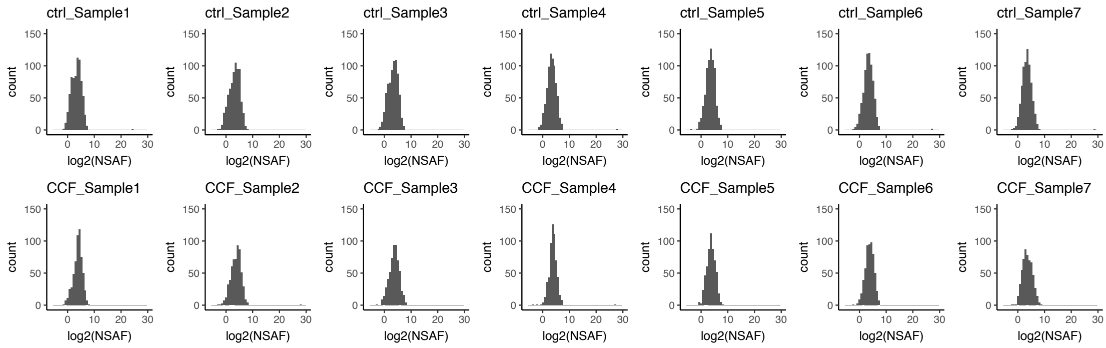


**Supplementary Figure S2:** The distribution of log2 transformed relative protein abundances (normalized spectral abundance factor (NSAF, materials and methods in main text)) approximates normal distribution in all samples and is similar between samples from the same patient. ctrl = control samples, CCF = clear cell foci samples, Sample1-Sample7 denote different patients.

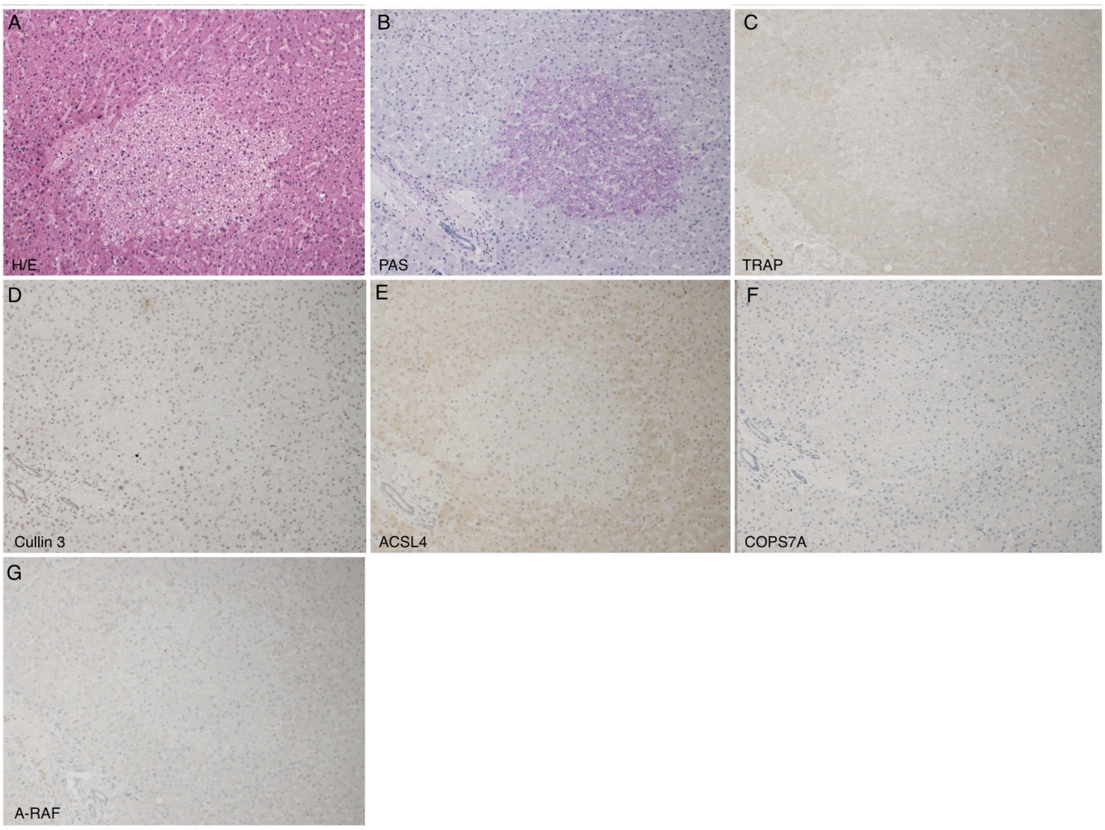


**Supplementary Figure S3:** Histochemistry and immunohistochemistry of representative human liver specimens with CCF and neighboring tissue. Hematoxilin/Eosin (H/E) staining (A), PAS reaction (B), and immunohistochemical detection of TRAP (C), Cullin 3 (D), ACSL4 (E), COPS7A (F), and A-RAF (G).

### SUPPLEMENTARY TABLES

**Table S1:** RNA concentration and quality as determined by Nanodrop and Bioanalyzer respectively

| Sample | RNA concentration (ng/µl) | RNA yield (ng) | RIN |
|---|---|---|---|
| sample 1 - H | 23 | 688 | 7,1 |
| sample 1 - N | 24 | 717 | 7,9 |
| sample 2 - H | 25 | 744 | 7,2 |
| sample 2 - N | 29 | 862 | 7,1 |
| sample 3 - H | 24 | 725 | 7,3 |
| sample 3 - N | 20 | 615 | 7,2 |
| sample 4 - H | 50 | 1514 | 6,9 |
| sample 4 - N | 36 | 1081 | 6,3 |
| sample 5 - H | 24 | 717 | 7,1 |
| sample 5 - N | 28 | 1120 | 7,0 |
| sample 6 - H | 17 | 520 | 6,9 |
| sample 6 - N | 23 | 705 | 6,9 |
| sample 7 - H | 26 | 765 | 7,2 |
| sample 7 - N | 26 | 793 | 7,2 |
| sample 8 - H | 20 | 609 | 6,4 |
| sample 8 - N | 22 | 645 | 6,4 |
| sample 9 - H | 16 | 470 | 6,8 |
| sample 9 - N | 25 | 738 | 6,3 |
| sample 10 - H | 44 | 1760 | 8,5 |
| sample 10 - N | 66 | 2640 | 6,2 |

**Table S2:** Antibodies used for immunohistochemistry

| Protein | product # / [reference] | company / source | antigen unmasking | antibody dilution |
|---|---|---|---|---|
| ACSL4 | HPA005552 | SigmaAldrich / Atlas Antibodies | boiling, pH 6 | 1:100 |
| Cullin 3 | 342200 | Life technologies | boiling, pH 6 | 1:100 |
| COPS7A | HPA026915 | SigmaAldrich / Atlas Antibodies | boiling, pH 6 | 1:100 |
| CYP5B | 15469-1-AP | ProteinTech | boiling, pH 6 | 1:100 |
| HSP10 | HPA038755 | SigmaAldrich /Atlas Antibodies | boiling, pH 6 | 1:1000 |
| LGAL7 | sc-271473 | Santa Cruz Biotechnology | boiling, pH 6 | 1:50 |
| monad/WDR92 | [1] | Makio Saeki<br>Division of Dental Pharmacology<br>Niigata University Graduate School of Medical and Dental Sciences<br>2-5274 Gakkocho-dori, Chuo-ku,<br>Niigata, 951-8514, Japan | boiling, pH 6 | 1:100 |
| RAB12 | HPA040727 | SigmaAldrich / Atlas Antibodies | boiling, pH 6 | 1:100 |
| RAB35 | HPA054146 | SigmaAldrich / Atlas Antibodies | boiling, pH 6 | 1:100 |
| A-RAF | HPA066326 | SigmaAldrich /Atlas Antibodies | boiling, pH 6 | 1:100 |
| STBD1 | HPA011952 | SigmaAldrich / Atlas Antibodies | boiling, pH 6 | 1:100 |
| TRAP 1 | HPA041082 | SigmaAldrich / Atlas Antibodies | boiling, pH 6 | 1:100 |
| USP28 | HPA006778 | SigmaAldrich / Atlas Antibodies | boiling, pH 6 | 1: 50 |

### Literature references


1. Saeki M, Irie Y, Ni L, Yoshida M, Itsuki Y, Kamisaki Y. Monad, a WD40 repeat protein, promotes apoptosis induced by TNF-alpha. Biochem Biophys Res Commun. 2006; 342: 568–72. doi: 10.1016/j.bbrc.2006.02.009.
